## Supplemental Figures for "A high-throughput yeast display approach to profile pathogen proteomes for MHC-II binding"

**Supplemental Information**

Supplemental Figures 1-7 (this document)

Supplemental Data (provided .xlsx file). Peptides and read counts for defined libraries and inferred registers for enriched peptides.

| | | Doped | | | | | Undoped | | | | | | |
|---|---|---|---|---|---|---|---|---|---|---|---|---|---|
| | | R0 | R1 | R2 | R3 | R4 | R0 | R1+ | R1- | R2+ | R2- | R3+ | R3- |
| Doped | R0 | 1.00 | 0.16 | 0.16 | 0.16 | 0.16 | 0.09 | 0.18 | -0.03 | 0.18 | 0.11 | 0.16 | 0.16 |
| | R1 | 0.16 | 1.00 | 0.91 | 0.86 | 0.75 | 0.24 | 0.76 | -0.30 | 0.78 | 0.42 | 0.72 | 0.72 |
| | R2 | 0.16 | 0.91 | 1.00 | 0.95 | 0.85 | 0.21 | 0.79 | -0.35 | 0.86 | 0.37 | 0.81 | 0.76 |
| | R3 | 0.16 | 0.86 | 0.95 | 1.00 | 0.94 | 0.17 | 0.75 | -0.37 | 0.89 | 0.25 | 0.88 | 0.69 |
| | R4 | 0.16 | 0.75 | 0.85 | 0.94 | 1.00 | 0.13 | 0.65 | -0.34 | 0.84 | 0.11 | 0.90 | 0.53 |
| Undoped | R0 | 0.09 | 0.24 | 0.21 | 0.17 | 0.13 | 1.00 | 0.36 | 0.41 | 0.21 | 0.42 | 0.16 | 0.27 |
| | R1+ | 0.18 | 0.76 | 0.79 | 0.75 | 0.65 | 0.36 | 1.00 | -0.27 | 0.86 | 0.66 | 0.76 | 0.87 |
| | R1- | -0.03 | -0.30 | -0.35 | -0.37 | -0.34 | 0.41 | -0.27 | 1.00 | -0.40 | 0.04 | -0.38 | -0.32 |
| | R2+ | 0.18 | 0.78 | 0.86 | 0.89 | 0.84 | 0.21 | 0.86 | -0.40 | 1.00 | 0.34 | 0.94 | 0.81 |
| | R2- | 0.11 | 0.42 | 0.37 | 0.25 | 0.11 | 0.42 | 0.66 | 0.04 | 0.34 | 1.00 | 0.18 | 0.62 |
| | R3+ | 0.16 | 0.72 | 0.81 | 0.88 | 0.90 | 0.16 | 0.76 | -0.38 | 0.94 | 0.18 | 1.00 | 0.66 |
| | R3- | 0.16 | 0.72 | 0.76 | 0.69 | 0.53 | 0.27 | 0.87 | -0.32 | 0.81 | 0.62 | 0.66 | 1.00 |

**Supplemental Figure 1. Correlations between selection rounds.** Pearson correlation for HLA-DR401 SARS-CoV-2 and SARS-CoV defined library members (+/- signs indicate enriched (+) or not enriched (-) yeast in undoped library; rounds of selection are indicated e.g. “R1” indicates “Round 1”, and “R0” is the unselected “Round 0” library).

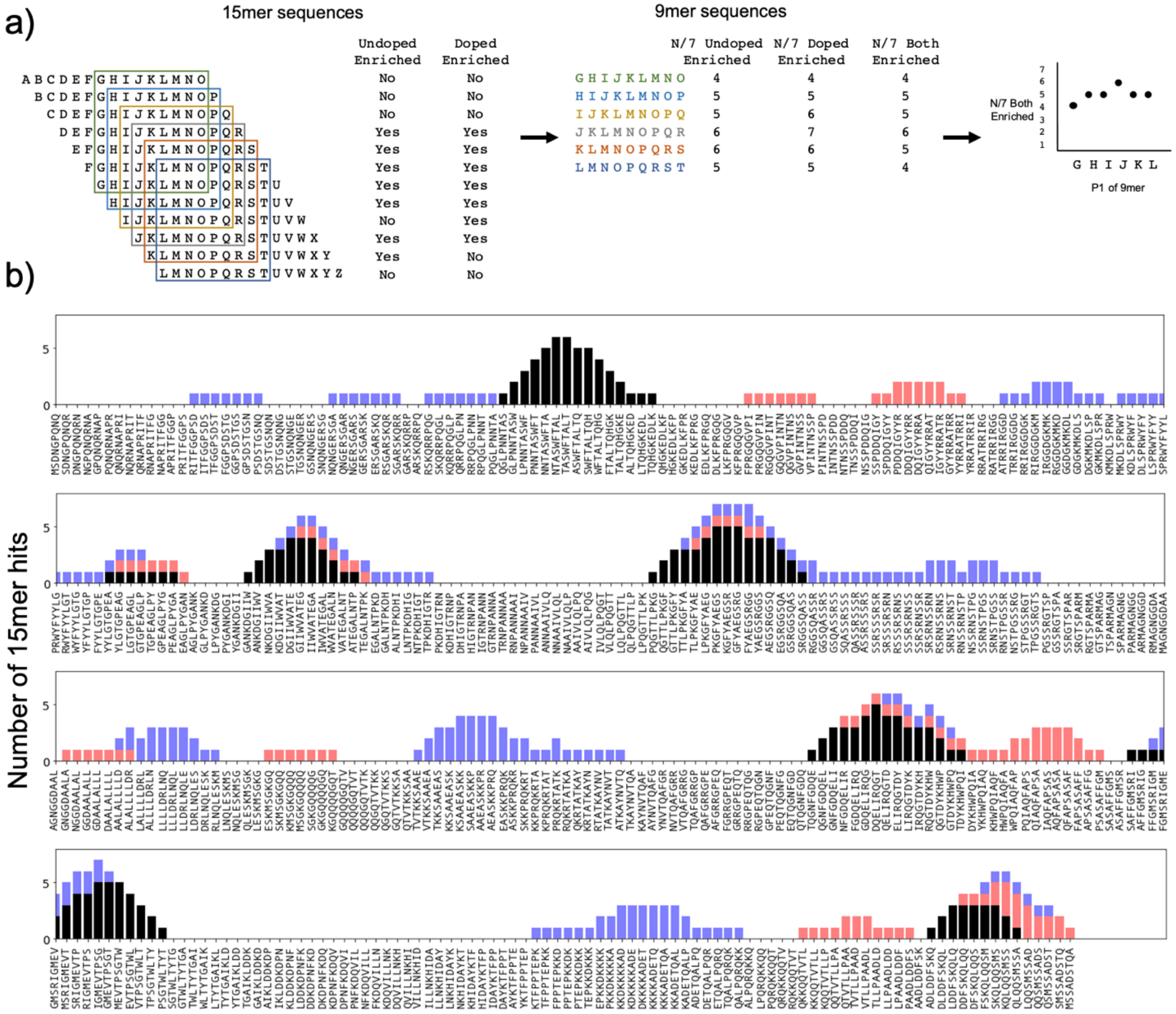


**Supplemental Figure 2. Utilizing overlapping peptides to call high confidence peptides.**
**a)** Schematic showing use of overlapping 15mers, containing redundant 9mers. Each 9mer is present in seven 15mers; for each 9mer, we calculate how many of these seven 15mers enriched. **b)** Number of peptides containing a given 9mer that are hits, for SARS-CoV-2 nucleocapsid on HLA-DR401. Black=hits in both undoped and doped libraries; blue=hits in undoped library only; red=hits in doped library only. Enrichment categories are stacked, for a maximum of seven 15mer hits, since each 9mer is present in seven 15mers.

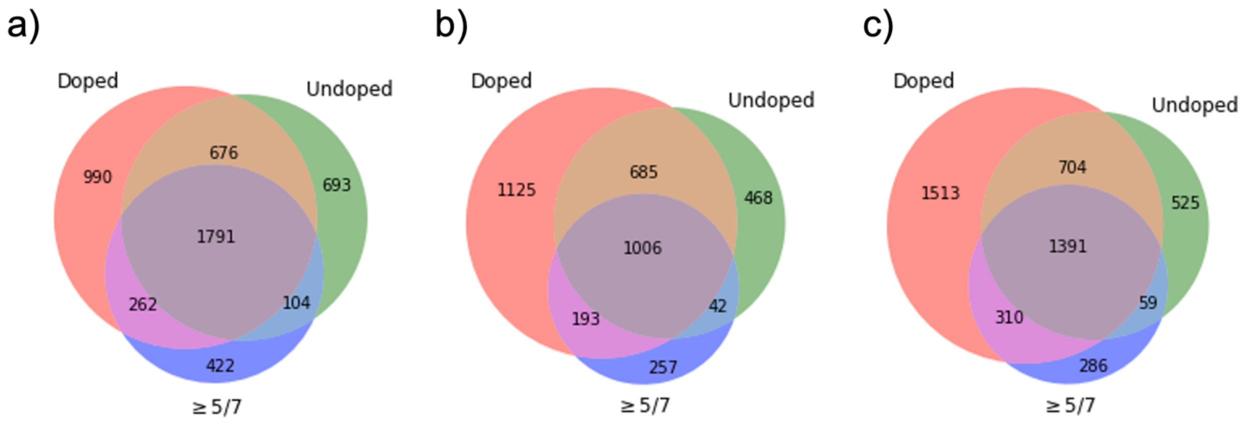


**Supplemental Figure 3. Full Venn diagrams.** Full Venn diagrams showing relationships between peptides which enriched in the doped library ("Doped"), and undoped library ("Undoped"), and contained a 9mer peptide which enriched in five or more of the seven 15mers containing it ("≥5/7"), for **a)** HLA-DR401, **b)** HLA-DR402, and **c)** HLA-DR404.

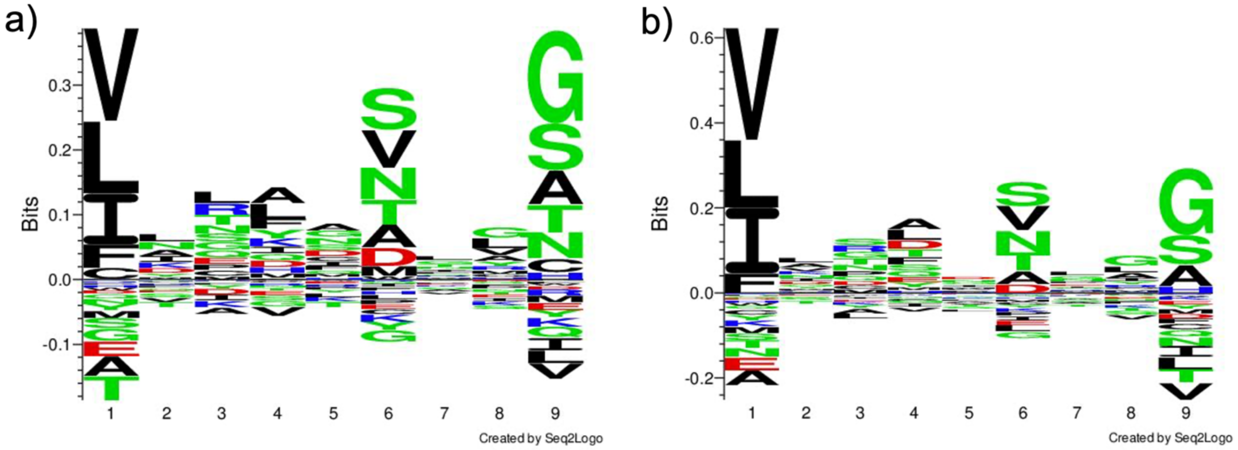


**Supplemental Figure 4. Sequence logo for HLA-DR402 and HLA-DR404. a)** HLA-DR402: Sequence logo of 1,690 peptides that enriched in both doped and undoped selections of the SARS-CoV and SARS-CoV-2 library for HLA-DR402. **b)** HLA-DR404: Sequence logo of 2,094 peptides that enriched in both doped and undoped selections of the SARS-CoV and SARS-CoV-2 library for HLA-DR404. Logos were generated with Seq2Logo-2.0.

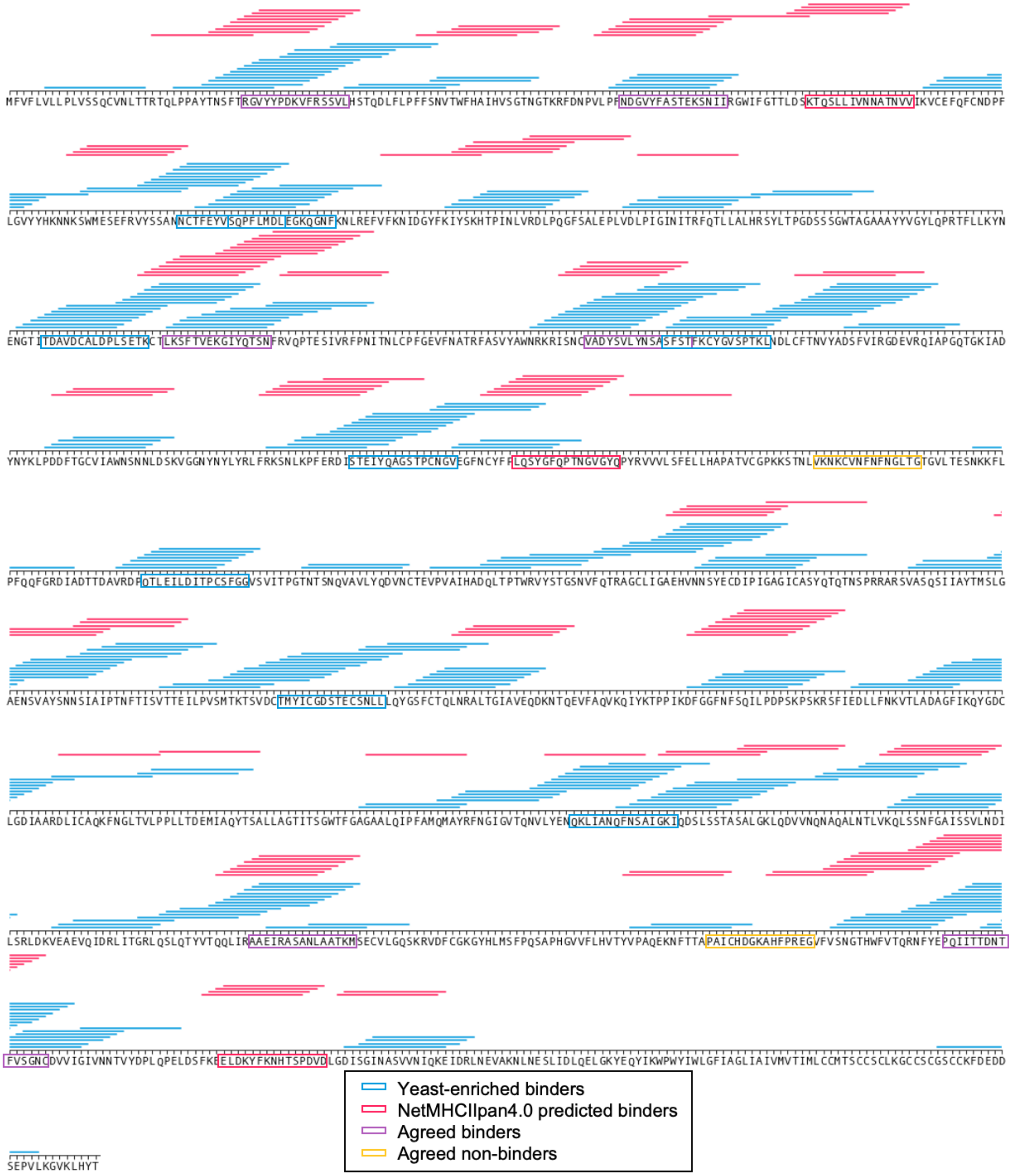


**Supplemental Figure 5. Comparing defined library selection with algorithmic predictions: SARS-CoV-2 spike protein.** 15mer peptides which enriched for binding to HLA-DR401 in both the doped and undoped libraries are indicated with horizontal lines above the enriched 15mer sequence (blue). NetMHCIIpan4.0 predicted binders (rank ≤ 10%) on yeast-formatted peptides are shown in red. Boxed sequences are tested in subsequent fluorescent polarization experiments, and colored as indicated in the legend.

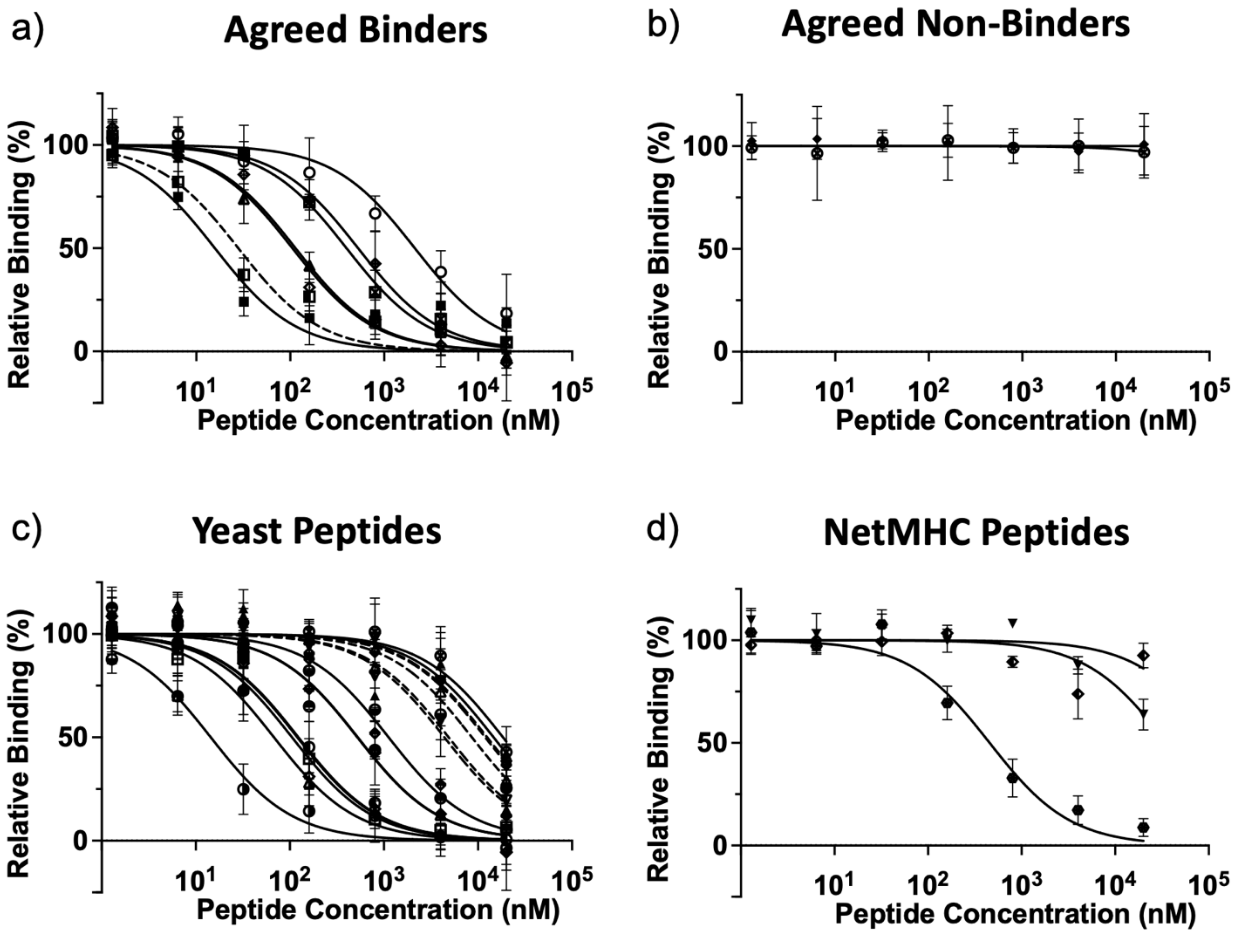


**Supplemental Figure 6. Titration curves for peptides tested via fluorescence polarization for binding to HLA-DR401, by category. a)** Agreed binder peptides which are predicted to bind by NetMHCIIpan4.0 and enriched in yeast display experiments. Dashed line is the positive control HA peptide. **b)** Agreed non-binder peptides which did not enrich in yeast display experiments and were not predicted to bind by NetMHCIIpan4.0. **c)** Yeast enriched peptides from Table 2 and Table 3. Offset variants from Table 3 are dashed lines. **d)** NetMHCIIpan4.0 predicted peptides which are not enriched in the yeast display library.

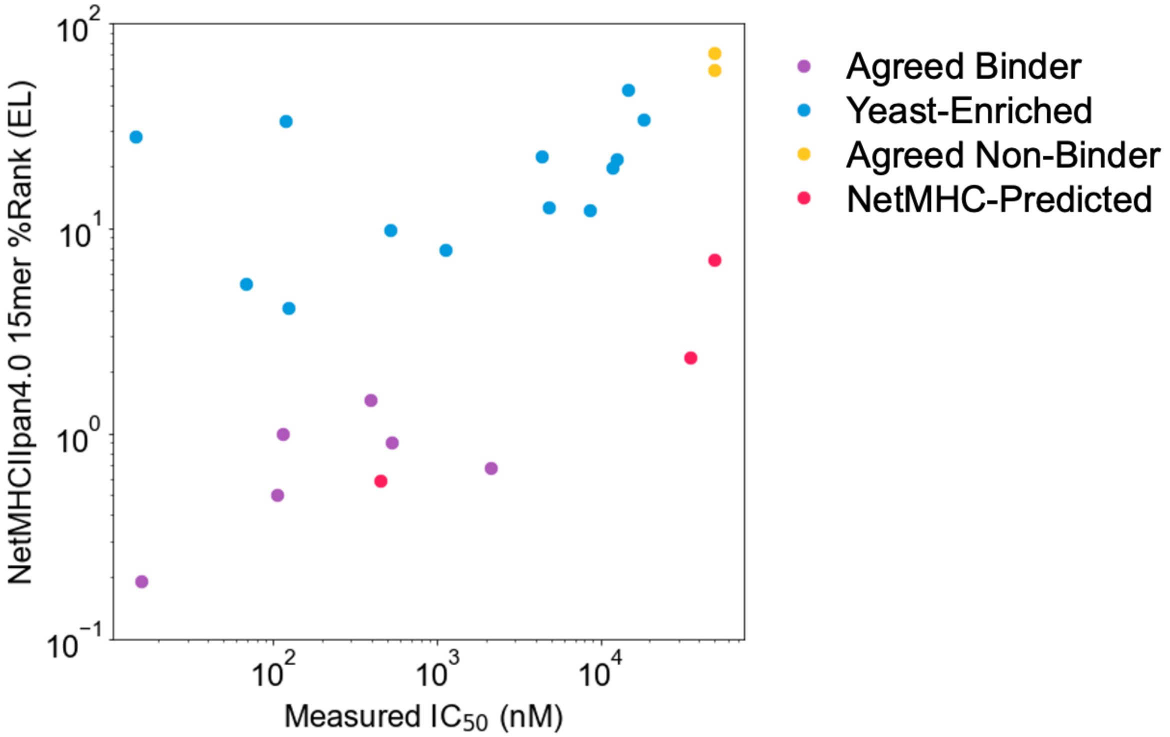


**Supplemental Figure 7. Comparing measured $IC_{50}$ values and prediction.** Relationship between measured $IC_{50}$ values and NetMHCIIpan4.0 predicted ranks in Eluted Ligand mode (EL) on unflanked (native) 15mer sequences. Data points are colored by label, and $IC_{50}$ values ≥50 µM are set to 50 µM.
